# Accelerating manufacturing to enable large-scale supply of a new adenovirus-vectored vaccine within 100 days

**DOI:** 10.1101/2021.12.22.473478

**Authors:** Carina C D Joe, Rameswara R Segireddy, Cathy Oliveira, Adam Berg, Yuanyuan Li, Dimitrios Doultsinos, Nitin Chopra, Steffi Scholze, Asma Ahmad, Piergiuseppe Nestola, Julia Niemann, Alexander D Douglas

**Author notes:** **Correspondence**, Alexander D Douglas, Jenner Institute, Henry Wellcome Building of Genomic Medicine, Old Road Campus, Oxford, OX3 7BN. **Funding information**, This work was funded by AstraZeneca and the EPSRC (EP/R013756/1; The Future Vaccine Manufacturing Research Hub), and benefited from in-kind support from Repligen Corporation and Sartorius. AstraZeneca reviewed the final manuscript for accuracy.

## Abstract

The Coalition for Epidemic Preparedness Innovations’ ‘100-day moonshot’ aspires to launch a new vaccine within 100 days of pathogen identification. Here, we describe work to optimize adenovirus vector manufacturing for rapid response, by minimizing time to clinical trial and first large-scale supply, and maximizing the output from the available manufacturing footprint.

We describe a rapid viral seed expansion workflow that allows vaccine release to clinical trials within 60 days of antigen sequence identification, followed by vaccine release from globally distributed sites within a further 40 days. We also describe a new perfusion-based upstream production process, designed to maximize output while retaining simplicity and suitability for existing manufacturing facilities. This improves upstream volumetric productivity of ChAdOx1 nCoV-19 by around four-fold and remains compatible with the existing downstream process, yielding drug substance sufficient for 10000 doses from each liter of bioreactor capacity.

Transition to a new production process across a large manufacturing network is a major task. In the short term, the rapid seed generation workflow could be used with the existing production process. We also use techno-economic modelling to show that, if linear scale-up were achieved, a single cleanroom containing two 2000 L bioreactors running our new perfusion-based process could supply bulk drug substance for around 120 million doses each month, costing <0.20 EUR/dose. We estimate that a manufacturing network with 32000 L of bioreactor capacity could release around 1 billion doses of a new vaccine within 130 days of genomic sequencing of a new pathogen, in a hypothetical ‘surge campaign’ with suitable prior preparation and resources, including adequate fill-and-finish capacity.

This accelerated manufacturing process, along with other advantages such as thermal stability, supports the ongoing value of adenovirus-vectored vaccines as a rapidly adaptable and deployable platform for emergency response.

## Introduction

The Coalition for Epidemic Preparedness Innovations and Pandemic Preparedness Partnership have proposed a ‘100 day mission’, or ‘moonshot’ aspiration of compressing the time taken for launch of a new vaccine to 100 days from pathogen identification.(1) Vaccine platform technologies with known safety, immunogenicity and manufacturing characteristics will be critical to reaching this goal, including mRNA and adenoviral vectors.

Adenovirus vectors offer a rapidly adaptable and deployable platform with proven safety and efficacy. Properties such as suitability for refrigerated storage, ability of a single dose to achieve protection lasting months, and widespread manufacturing technology transfer make the platform particularly attractive in low- and middle-income countries. More than two billion doses of the ‘Oxford/AstraZeneca’ adenovirus-vectored COVID-19 vaccine ChAdOx1 nCoV-19 (AZD1222, Vaxzevria) have been supplied. The product has been used in 177 countries worldwide and manufacturing is ongoing in 15 countries across five continents (2, 3). This was enabled by rapid transfer of the platform manufacturing process to multiple production facilities. Despite this success, manufacturing speed remains a potential limitation of adenoviral-vectored vaccines compared with other platforms including mRNA-based vaccines. We therefore aimed to accelerate adenovirus vector manufacturing further to maximize the suitability of this platform for ongoing and future responses to emerging pathogens and new variants.

Here, we describe work seeking to optimize each of three measures of a vaccine platform technology’s suitability for emergency response to emerging pathogens and variants: time from pathogen sequence identification to supply of vaccine to clinical trial; time to first large-scale product release; and time to release of one billion doses.

The two activities required for adenovirus manufacturing ‘start-up’ are the expansion of host cells and the generation of seed virus with which to infect those cells (assuming that facilities, equipment, staff and materials are already in place). These activities can take place in parallel, and the latter is slower, making Good Manufacturing Practice (GMP)-compliant virus seed generation the limiting factor for initiation of both trial-scale and large-scale manufacturing. This requires synthesis of DNA encoding the antigen, insertion of this transgene into the adenoviral genomic construct, transfection of the genomic construct into producer cells to ‘rescue’ the virus, limiting dilution to isolate clonal virus, and serial amplification. GMP requires time-consuming quality-control assays at multiple points in the seed-production process, but the probability of failure is low, in our experience, for the most time-consuming assays (detection of adventitious microbes and confirmation of genetic stability). Seed generation time can therefore be reduced by proceeding ‘at financial risk’ to subsequent steps, in advance of assay results, provided that all required results will be available before vaccine release.

Following manufacturing start-up, the ramp-up capacity is constrained by the volumetric productivity of the manufacturing process and the facility cycle time (interval between batches). These factors determine the number of doses which can be produced each month per liter of bioreactor capacity, or per square meter of facility footprint.

Current worldwide supply of ChAdOx1 nCoV-19 uses a process built upon a method that we developed before the COVID-19 pandemic to enable rapid responses to emerging pathogen outbreaks (4, 5). This was specifically designed to be quick and straightforward to adopt for manufacturing of any adenovirus-vectored vaccine in any suitable facility, including those in low- and middle-income countries. Use of a GMP-compliant bank of suspension T-REx-293 cells (ThermoFisher) enables repression of transgene expression during production, which overcomes the potential problem of viral growth impairment by the transgene product. Only single-use, off-the-shelf components are used throughout manufacturing, including both the upstream fed-batch stirred-tank bioreactor process and the downstream tangential-flow filtration and membrane-based anion-exchange chromatography process.

The volumetric productivity of the current ChAdOx1 nCoV-19 production process is approximately 2 × 10^11^ VP/mL of bioreactor culture, or approximately 2500 final doses/L (5). Producing a final dose of 5 × 10^10^ VP requires upstream production of approximately 1.5 × 10^11^ VP because of losses during purification of drug substance (DS; approximately 50% loss), fill-finish and extraction from vials (a further 33% of the drug substance, approximately), but relatively little scope exists for improving the efficiency of these manufacturing stages. Major output improvement thus requires improved upstream volumetric productivity. Volumetric productivity (VP/mL) is the viable cell density (cells/mL) multiplied by the cell-specific productivity (VP/cell) during viral vector replication in the producer cell line (i.e. volumetric productivity = viable cell density × cell-specific productivity).

The ‘cell density effect’ is the principal factor limiting volumetric productivity of adenoviral vectors (6, 7). Cell densities above 10^7^ viable cells/mL are readily achievable in fed-batch growth of uninfected producer cell lines in stirred-tank bioreactors (8). Unfortunately, cell-specific productivity of adenoviral vectors drops very sharply at quite modest cell densities, above approximately 1–2 × 10^6^ viable cells/mL at infection (corresponding to somewhat higher peak viable cell density after infection) (9). As a result, increasing the cell density beyond this modest level does not improve volumetric productivity of adenoviral vectors (6, 7, 10). High cell-specific productivity has an important additional benefit on top of its effect upon volumetric productivity, in that a high ratio of active product to cell-derived impurities facilitates downstream processing.

In our fed-batch ChAdOx1 nCoV-19 manufacturing process at large scale, we have observed cell-specific productivity of approximately 3 to 5 × 10^4^ VP/cell, with a peak viable cell density in the region of 4 to 6 × 10^6^ cells/mL (5). This cell specific productivity value is in line with previously published results with ChAdOx1 and other species E simian adenovirus vectors and with human adenovirus vectors (4, 7). The combination of cells, antigen repression, vector engineering, medium and feed allows this cell-specific productivity to be maintained at somewhat higher cell densities than previously reported, resulting in increased volumetric productivity. Nonetheless, overcoming the cell density effect remains a central challenge in boosting ChAdOx1 nCoV-19 manufacturing output (5).

The physiological basis of the cell density effect is incompletely understood, but an inhibitory effect of cellular waste products upon viral replication is a likely contributor. Production of adenovirus vectors is fundamentally a batch process because a single cycle of viral replication lyses the producer cells over a time-frame of 24 to 72 hours (7). This contrasts with production of other biological products like monoclonal antibodies, where constitutive expression enables continuous processes. Although viral replication rapidly increases cellular energy demand (11), nutrient supplementation with glucose, vitamins, amino acids or nucleotides does not raise the cell density effect barrier beyond about 1–2 × 10^6^ viable cells/mL at infection (6, 9, 11). Cell-specific productivity is impaired by metabolic waste products such as ammonia (from glutamine metabolism) and lactate (from anaerobic glucose metabolism) (11–13). Medium exchange by centrifugation or tangential flow filtration before or during the brief period of adenoviral vector replication in the producer cells can both replenish nutrients and remove waste products. Alternating tangential-flow filtration (ATF) and levitating magnetic pump head systems can accomplish this with low shear forces, an important consideration for fragile virus-infected producer cells. In contrast to nutrient supplementation, medium exchange by perfusion is reportedly effective in raising the cell density effect barrier in processes using species C human adenovirus vectors (14–17). As compared to fed batch processes, however, perfusion requires additional equipment and the use and handling of substantial quantities of additional medium. It is perceived by some as being a complex operation which some manufacturing sites may find difficult to implement.

Here, we demonstrate that medium exchange using ATF can raise the cell density effect barrier for production of ChAdOx1 nCov-19. Using the simplest possible perfusion conditions, we show that increasing viable cell density at infection to 6 × 10^6^ cells/mL improves volumetric productivity by approximately four fold, with no loss of cell-specific productivity. We also provide a workflow for rapid production of working virus seed to support our new process at global scale, enabling large-scale vaccine release within 100 days from pathogen sequence identification. Using a techno-economic model of the new method at large scale, we estimate the output from a typical facility, cost of goods, and the facility scale needed to produce 1 billion doses of a future vaccine within 130 days of genomic sequencing of a novel pathogen.

## Methods

### CELLS AND VACCINE

All experiments used a previously described research cell bank of T-REx-293 human embryonic kidney (HEK) cells stably expressing the tetracycline repressor protein (ThermoFisher) (4). Cells were adapted to antibiotic-free and serum-free BalanCD HEK293 medium (Irvine) and maintained as previously described (4). Cells were expanded in medium supplemented with 4 mM L-alanyl-L-glutamine (Gibco) in 125 mL and 1 L Erlenmeyer flasks (Corning) and 5 L Optimum Growth (Thomson) flasks using humidified shaking incubators (Kühner) at 37°C, 8% CO2 and 130 rpm with a 25 mm orbit. Cells were sub-cultured every 3 to 4 days to maintain viable cell density between 0.5 and 3 × 10^6^ cells/mL at a maximum fill volume of 20%. Viable cell density was measured daily using a NucleoCounter NC-202 (ChemoMetec).

ChAdOx1 nCoV-19 was derived as previously reported (18). A seed stock was prepared for the present study using our previously described fed-batch manufacturing process (4). Except where otherwise stated, seed was used to infect cells at an multiplicity of infection (MOI) of 5 IU/cell.

### RAPID SEED VIRUS PRODUCTION

An Erlenmeyer flask with a culture volume of 30 mL and a viable cell density of 0.8 × 10^6^ cells/mL received ChAdOx1 nCoV-19 at an MOI of 6 VP/cell (as determined by qPCR). This qPCR value corresponds to approximately 0.1 IU/cell with a plausible range of 0.025 to 0.2 IU/cell assuming a typical particle:infectivity ratio for ChAdOx1 nCoV-19. The cells were harvested and lysed 6 days later. Virus was partially purified using depth filtration and anion exchange chromatography with syringes and small-scale filters (C0SP 23 cm^2^ [Merck] and Sartobind Q Nano [Sartorius]). Buffer compositions and other parameters were as previously described but scaled proportionately to the product and filter sizes (5). Bioreactors were seeded with approximately 0.5 × 10^6^ cells/mL in approximately 80% of the maximum working volume, and infected 28–30 hours later at a MOI of 6 VP/cell and viable cell density of approximately 0.8 × 10^6^ cells/mL. Cell cultures were fed with BalanCD medium 5% v/v at 48 hours and 96 hours after infection (±4 hours). Temperature was reduced within 4 hours of the second feed. Cultures were harvested approximately 140 hours after infection and virus was partially purified as described above.

### HIGH-THROUGHPUT PERFUSION STUDIES

#### Multi-parallel bioreactors

For high-throughput screening of process conditions, the multi-parallel bioreactor and perfusion system Ambr^®^250 High Throughput Perfusion (Ambr250 HT Perfusion; Sartorius) was used with 0.2 μm hollow-fibre filters in ATF mode. The total reactor volume was 210 mL per vessel and the stirring power input was 40 W/m^3^. Temperature was controlled at 37°C, dissolved oxygen (DO) was controlled at 55 % by sparging with air and/or O2, and *p*H was controlled at 7.25 (dead-band, 0.05) by sparging with CO2 mix or addition of 1M sodium bicarbonate. Antifoam C emulsion 30% (Sigma-Aldrich) was diluted to 2% and automatically added as required via the Ambr250 integrated foam sensor and liquid handler. Each Ambr250 vessel was filled with pre-warmed BalanCD HEK293 medium and seeded with 0.5 × 10^6^ viable T-REx-293 cells/mL. After 3 days, cells were bled and replenished to reactor working volume containing 1 × 10^6^ viable cells/mL (thus modelling pre-production ‘n-1’ cell seed culture expansion in a stirred tank reactor). Nutrient supplementation used 5% BalanCD HEK293 feed, as described below.

Viable cell density was measured daily using a Cedex HiRes^®^ Analyzer (Roche). Metabolites were automatically analysed daily via the integrated BioProfile^®^ FLEX2 device (Nova Biomedical). Samples for adenoviral vector quantification by quantitative PCR (see below) were taken 44 and 48 hours after infection. For each timepoint, VP counts were calculated as the mean of triplicate samples, each analysed in triplicate wells on triplicate plates. The mean of the 44-hour and 48-hour values was used for analysis.

#### Experimental design

A ‘design of experiments’ approach was used to determine the influence of three factors on productivity: perfusion start time, at a rate of one reactor volume per day; high or low viable cell density at infection; and duration of intensified perfusion (at a rate of two reactor volumes per day, starting at infection). When the duration of intensified perfusion was zero, perfusion continued at one reactor volume per day after infection. Two successive full-factorial experiments each comprised two levels of these three factors (2 × 2 × 2), in eight single bioreactors. Each experiment also included one ‘centre-point’ condition (in triplicate bioreactors) and one ‘simplified centre-point-like’ condition (singly or in duplicate), for a total of 12 parallel bioreactors in the first experiment and 13 in the second (Supplementary Table 1).

The first experiment was designed to target a viable cell density at infection of approximately 4 or 12 × 10^6^ cells/mL with a center point of c. 8 × 10^6^ cells/mL; a perfusion start time of 12 or 48 hours before infection, with a center point of 24 hours; and an intensified perfusion duration of 0 or 48 hours after infection, with a center point of 24 hours. In the second experiment, the design was adjusted to target viable cell densities at infection of approximately 6 or 18 × 10^6^ cells/mL with a center point of c. 12 × 10^6^ cells/mL; the perfusion start time was set to 48 or 72 hours after cell density adjustment by reactor bleed, with a center point of 60 hours; and the duration of intensified perfusion was as in the first experiment.

The ‘simplified center-point-like’ condition used center-point viable cell density at infection, early perfusion start time and an intensified perfusion duration of 0 hours (i.e. no intensification of perfusion; continuous perfusion at one reactor volume per day). This condition was designed to balance high productivity (via early perfusion start and intermediate cell density) with suitability of the process for multiple facilities (avoiding the facility-fit and medium-supply challenges which might arise from handling two vessel volumes of medium per day at large scale).

The difference in triggering of perfusion start time between the two experiments provided a range of viable cell density at the time perfusion was started, with higher cell densities in the second experiment than the first. The experiments were performed successively, enabling observations made during the first experiment to inform design of the second. For the second experiment the maximum fill volume during the last shake flask cultivation step was reduced to 10% (to reduce aggregation of the cells before inoculation of the Ambr250 HT Perfusion bioreactor vessels). In addition, nutrient supplementation was increased by addition of 5% feed solution at the bioreactor bleed three days after inoculation (to support the higher cell densities in the second experiment).

Two response variables were defined: volumetric productivity (VP/mL), and cell-specific productivity (VP/cell). Cell-specific productivity values were based on the peak viable cell density for the reactor recorded by the Cedex analyzer.

#### Statistical analyses

Analyses were performed using MODDE version 13 (Sartorius). The main analysis combined the two experiments using a dummy variable denoting each experiment (EXP). To combine the experiments, the perfusion start-time factor (PST) was replaced with a continuous variable describing the perfusion-start viable cell density (PSVCD). Where necessary, PSVCD was calculated by extrapolation or interpolation from the nearest available cell count and population doubling time (in all cases, within 24 hours of perfusion start). One centre-point condition reactor from experiment 2 was excluded from all analyses due to microbial contamination. The response variables were normalized with a log_10_ transformation. In the main analysis, a model including the terms EXP, PSVCD, viable cell density at infection (VCDI) and intensified perfusion duration (IPD) was fit by partial least squares regression. Statistical significance of parameter coefficients was assessed using *t* tests (with a null hypothesis that parameter coefficients were zero) and values of *r^2^* and *q^2^* were calculated using functions in MODDE as defined in the software user guide.

Eight sensitivity analyses were performed to evaluate the impact on the overall conclusions of the following changes from the above analysis. First, the PST factor was replaced with perfusion duration before infection (PDBI) instead of PSVCD. Secondly, raw response data was used instead of log10-transformed data. Thirdly and fourthly, each experiment was analysed individually instead of in combination. Fifthly, the experiments were pooled without using the EXP dummy variable. Sixthly, three bioreactors with simplified centre-point-like conditions that did not fit the full-factorial design were excluded. Seventhly, two bioreactors were excluded due to initial low cell density likely resulting from error during set-up (one centre-point reactor in the first experiment and one simplified centre-point-like reactor in the second experiment). Finally, the model was fitted using multiple linear regression instead of partial least-squares regression.

### SCALED-DOWN PRODUCTION BIOREACTORS

We designed perfusion filter dimension and flow parameters, using Repligen XCell Lab technology to provide an accurate 3L scaled-down model of production at 2000L scale. A 0.022 m^2^ filter (ATF1, Repligen) used with alternating tangential flow at 5 L/m^2^/min to exchange medium at 3 L/day (0.11 L/m^2^/min) provides shear rate (1516 s^−1^) and other fluid dynamic parameters similar to those of an 11 m^2^ filter (ATF10, Repligen) used for perfusion of a 2000 L bioreactor of circa 1500 L working volume under recommended conditions. These flow rates fall comfortably within commonly used operating ranges.

BioBLU 3c single-use bioreactors with an open pipe, a pitched-blade impeller, and an optical *p*H port were used in a BioFLo320 parallel bioreactor system (Eppendorf) with integrated intelligent sensor management (Mettler Toledo). The submerged addition line was welded to the bioreactor connection line of an XCell ATF 1 single-use hollow-fibre filter and pump device (Repligen) with a 0.22 μm pore size, 1 mm lumen and a 218 cm^2^ nominal surface area. Alternating tangential flow was set to 0.11 L/min (5 L/m^2^/min) using an XCell Lab Controller (Repligen). The desired rate of medium exchange was achieved by controlling the flow rate of the permeate (spent medium) and adding new medium, both via the BioFlo320 integrated pumps (after calibration). The new medium feed pump was controlled by a gravimetric feedback loop programmed to maintain constant reactor weight.

Bioreactor growth medium was BalanCD HEK293 supplemented with 4 mM L-alanyl-L-glutamine, 0.01% antifoam C emulsion (Sigma-Aldrich) and 10–20% BalanCD feed (Irvine). Bioreactors were stirred at 160–180 rpm (26–33 W/m^3^) and maintained at 37°C using a heating jacket. Dissolved oxygen was controlled at >55% by sparging with air and/or O_2_ via a cascading control loop. The *p*H was controlled at 7.30 (dead-band, 0.1) by sparging with CO_2_ or addition of 7.5% sodium bicarbonate. Initial T-REx-293 cell density was 0.5 to 2 × 10^6^ viable cells/mL in a working volume of or 3 L. After infection with ChAdOx1 nCoV-19, samples were taken daily for adenoviral vector quantification and biochemical analysis using a Stat Profile Prime analyser (Nova Biomedical).

### HARVEST AND CONCENTRATION

Cells were lysed 42–48 hours after infection with ChAdOx1 nCoV-19 by addition of 1/9 culture volume of 10% v/v polysorbate 20, 50% w/v sucrose and 20 mM MgCl_2_ in 500 mM tris *p*H 8.0, 15 minutes after addition of benzonase to a final concentration of 100 U/mL (Merck Millipore). Dissolved oxygen and pH were uncontrolled during lysis but stirring and temperature control continued. After two hours, the lysate was clarified over 270 cm^2^ Millistak+ HC Pro C0SP depth filters coupled with Opticap XL150 0.2 μm sterile filters at a flow rate of 3.3 L/min/m^2^ as previously described (4). Turbidity was measured using a TN-100 waterproof turbidimeter (Thermo Scientific Eutech).

In one experiment, the clarified lysate was concentrated by tangential flow filtration at 5 L/min/m^2^ using a KR2i KrosFlo (Repligen) with Pellicon 2 mini cassettes (300 kDa, 0.1 m^2^, BioMax polyethersulfone membrane with C-screen; Merck), as previously described (4), but with the addition of a KRJr permeate control pump (Repligen) set to 0.66 L/min/m^2^. Feed pressure was controlled at 0.7 bar by an automated backpressure valve on the retentate outflow and diafiltration buffer feed rate was matched to the permeate flow rate by controlling the weight of the process reservoir. After 2-fold concentration, the retentate underwent diafiltration with one diavolume of ion exchange wash buffer (222mM NaCl, 1mM MgCl2 and 5% w/v sucrose in 50mM TrisHCl, *p*H 8.0, with a conductivity of 23–24 mS/cm).

### PURIFICATION

After clarification, lysates were purified by ion-exchange chromatography. AEX was preceded in one case (experiment CJ74) by limited tangential flow filtration as described above, and in others by dilution of lysate 1:3 with wash buffer (as above). Before chromatography, the conductivity of the lysate was adjusted to 23–24 mS/cm using 5M NaCl. Fresh single-use SartobindQ capsules with 150 mL bed volume and 8 mm bed height were used in a peristaltic pump-driven rig incorporating single-use ultraviolet absorbance, conductivity and pressure sensors (Pendotech), as previously described (4). Capsules were washed, equilibrated, loaded and eluted in accordance with the manufacturers instructions. Briefly, the membrane was sanitized with 30 membrane volumes (MV) of 1M NaOH at 1 MV/minute, activated with 10 MV of 1M NaCl at 2 MV/minute, and equilibrated with 20 MV of ion exchange wash buffer (see above) at 2 MV/minute. Samples were applied at 2.5–3.0 MV/minute, washed as per the equilibration step, and eluted with 5 MV of 444 mM NaCl, 1mM MgCl2 and 5% w/v sucrose in 50mM Tris HCl, *p*H 8.0, with a conductivity of 39–40 mS/cm, at 2 MV/minute. Eluate was collected into a reservoir containing a ‘cushion’ of charge buffer (35mM NaCl, 10mM histidine, 1mM MgCl2, 0.1 mM EDTA, 7.5% w/v sucrose, 0.5% v/v ethanol, *p*H 6.6).

After ion-exchange chromatography, the eluate buffer was exchanged with 6 diavolumes of formulation buffer A438 without polysorbate 80. This final tangential flow filtration step was performed using Pellicon 2 mini cassettes (as above) with a feed flow rate of 5 L/min/m^2^, a feed pressure of 0.7 bar, and a permeate flow rate of 0.66 L/min/m^2^ (matched by the rate of addition of fresh A438 via an automated auxiliary pump). Finally, polysorbate 80 concentration was added to a final concentration of 0.1% v/v and the product was passed through 0.2 μm filters (Nalgene).

### QUANTIFICATION AND QUALITY ASSESSMENT

ChAdOx1 nCoV-19 was quantified as previously described, using quantitative PCR (qPCR) and ultraviolet spectrophotometry assays for viral particles and a cell-based immunostaining assay for infectious units (4). ChAdOx1 nCoV-19 in cell-culture samples was quantified using qPCR and infectivity assays following cell lysis, essentially as described above for harvesting. ChAdOx1 nCoV-19 in ion-exchange eluate samples and in purified product was quantified by ultraviolet spectrophotometry before addition of polysorbate 80.

To improve qPCR accuracy, each sample was assayed in triplicate wells on triplicate plates with standard curve samples (10^6^ to 10^10^ genome copies spaced at tenfold intervals) and two internal controls. These comprised research-grade ChAdOx1 nCoV-19 purified by CsCl gradient ultracentrifugation and quantified by spectrophotometry, and clinical-grade ChAdOx1 nCoV-19 quantified by a GMP-qualified external qPCR assay (Advent, IRBM SpA, Italy). The acceptance criteria for qPCR values were: accuracy of internal controls within 15%; efficiency of 90–110%, and standard curve linearity (R^2^) of at least 0.99.

Residual host-cell protein (HCP) was quantified using the HEK293 HCP ELISA kit (Cygnus Technologies) according to the manufacturer’s instructions. Residual host-cell DNA was quantified using the previously reported qPCR method, with a lower limit of quantification of 100 pg/mL for intact HEK293 cell DNA (4). Polyacrylamide gel electrophoresis was conducted using standard molecular biology laboratory techniques.

### ELECTRON MICROSCOPY

Samples were diluted to 3–5 × 10^11^ VP/mL in water as required and applied to freshly glow discharged, carbon filmed, 300-mesh copper grids (TAAB Laboratories). Grids were incubated at room temperature for 2 minutes, blotted, immediately transferred to a 20 μL droplet of 2% uranyl acetate for 10 s, then blotted and air dried. Images were acquired at 120 keV on a Tecnai 12 transmission electron microscope (ThermoFisher) equipped with a OneView camera (Gatan).

### TECHNO-ECONOMIC MODELLING

Models were constructed and evaluated using Biosolve software (Biopharm Services). As a starting point, we used our previously described model of a 2000 L fed batch process (5). We made the following minor modifications to this base model based on subsequent experience: durations of each step in the seed train were revised to reflect a 36h cell population doubling time, and overall recovery in the downstream process was reduced from 65% to 50%. To model a 2000 L perfusion process, we examined two seed-train and production bioreactor configurations. First, we considered minimal change from the fed-batch seed train, with seeding of the production bioreactor from a 200 L ‘n–1’ bioreactor, no use of perfusion, and 7 days of cell expansion in a single production vessel to reach a density of approximately 6 × 10^6^ cells/mL before infection. Secondly we considered using a 200 L ‘n–2’ bioreactor to seed a 200 L perfusion-capable ‘n–1’ bioreactor for further cell expansion (reducing the time spent by each batch in the production vessel by 72h), with the single ‘n–1’ reactor servicing two 2000 L production bioreactors alternately.

At the harvest stage, the benzonase concentration was increased from 15 units/mL in the fed batch process model to 100 units/mL in the perfusion model. Filter loadings in the perfusion downstream process model were based upon those used in the fed-batch downstream process, assuming a 4-fold increase in both product and host cell material for a given volume of upstream culture. Volumetric loadings of the clarification, pre-anion-exchange filter (0.8/0.2 μm) anion-exchange filter, tangential-flow filter and bioburden reduction filter were thus all reduced to 25% of those used in the fed-batch model. Concentration in the tangential-flow filtration stage was limited to 0.33 x the bioreactor volume, corresponding to an expected product concentration of 2–3 × 10^12^ VP/mL. All these values were in line with our experimental experience.

## Results

### RAPID LARGE-SCALE VIRAL SEED SUPPLY

Most adenovirus production methods involve infection of all producer cells in a ‘single hit’ with an MOI above 1 IU/cell, followed by harvest approximately two days later, after a single viral lifecycle. Our previous work has used such processes throughout seed amplification. After limiting dilution to isolate clonal virus, we have typically used adherent cells to perform approximately 4 amplification steps in a pre-GMP laboratory. At each stage, virus is recovered by freeze-thaw-mediated cell lysis and the infectious titer in the lysate is measured to determine the MOI to use for the following step. A large ‘safety factor’ is typically allowed at each stage (i.e. oversizing to allow for the possibility of poor output). GMP master virus seed and working virus seed production then proceeds similarly, but in stirred tank reactors and using more efficient detergent-mediated lysis. It would be challenging to produce sufficient working virus seed using this approach to meet the needs of a global campaign where MOI is above 1 IU/cell in the manufacturing process. This was one of the factors which motivated development of the novel ‘two viral lifecycle’ process currently used for ChAdOx1 nCoV-19. Use of an MOI considerably less than 1 IU/cell in effect allows an additional cycle of viral replication to occur within the production bioreactor, and substantially reduces the requirement for input working virus seed. To summarize, our current approach thus involves high-MOI processes for seed generation, and a low-MOI process for drug substance production.

Here, we reversed the above approach, reasoning that the high viral ‘amplification factor’ achieved by a two-cycle / low-MOI culture would be ideal for generation of seed with the minimum number of amplification steps, and hence the minimum burden of testing between stages. By facilitating production of much larger quantities of working virus seed than previously, this new approach also enables a high-MOI production process.

Because low-MOI processes have now been extensively characterized in suspension cells, including the use of highly efficient detergent-mediated (rather than freeze-thaw) lysis to recover virus, we sought to make the earliest possible transition to suspension cell culture. To minimize delays for testing, we made the assumption that virus grown in cells from a previously tested master cell bank by skilled operators is highly unlikely to be contaminated with bacteria or mycoplasma. This allows immediate transfer of working virus seed to new facilities with acceptable financial risk (or on the basis of rapid assays). To further streamline testing between amplification steps, we reasoned that calculation of MOI on the basis of genome-containing viral particles would be an accurate enough estimate of infectious titre, given that particle:infectivity ratios are rarely outside the range 30 to 300. This approach would enable rapid qPCR assay of the input virus, rather than a time-consuming infectivity assay. Incorporation of depth filter clarification, anion exchange and 0.2 μm filtration, using appropriately scaled versions of the established downstream unit operations, would provide low bioburden seed virus in a controlled buffer matrix without detergent.

We designed a seed amplification workflow based upon these principles. We estimated that a single uncontrolled-MOI amplification step in adherent cells in a 6-well plate, followed by a low-MOI suspension-cell-based amplification step in a 30 mL shake flask would provide sufficient pre-master seed to infect a 50L stirred-tank bioreactor. This in turn could provide both ≥10 000 doses of clinical trial drug substance and adequate master virus seed to meet the needs of a global production campaign (Figure 1, Table 1).

**Figure 1.**
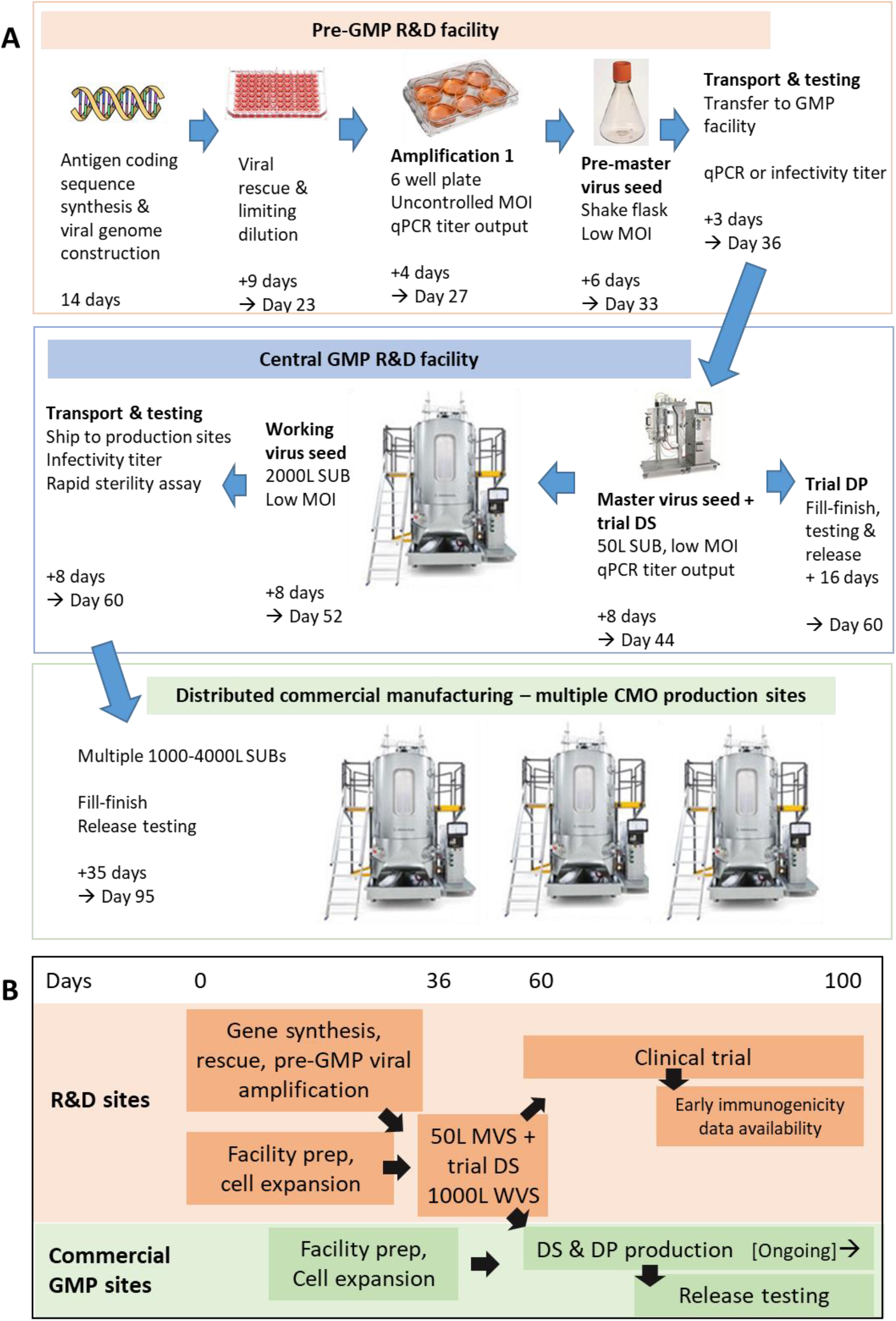
Rapid virus seed production enables early vaccine release. **(a)** Detailed scheme for virus seed production, including anticipated timing of each step. This timing assumes availability of facilities, equipment, materials and staff. It further assumes cell expansion for MVS, WVS and DS production occurring in parallel with preceding steps of virus seed generation, potentially making use of a ‘bleed and dilute’ strategy to hold cells at 50-200L volume and hence ensure immediate readiness for WVS / DS production upon seed availability. Finally, it assumes decisions to proceed ‘at financial risk’ at all stages, ahead of the availability of any test results other than those stated, but with completion of full testing before drug product release. **(b)** High level overview of development campaign combining seed production, supply of vaccine to clinical trial, execution of clinical trial, and large-scale manufacturing, enabling large-scale product release at day 100. Abbreviations: MVS, master virus seed; WVS, working virus seed; VP, viral particles; MOI, multiplicity of infection; GMP, good manufacturing practice; CMO, contract manufacturing organization; DS, drug substance; DP, drug product; USP, upstream process; SUB, single-use bioreactor.

**Table 1.**
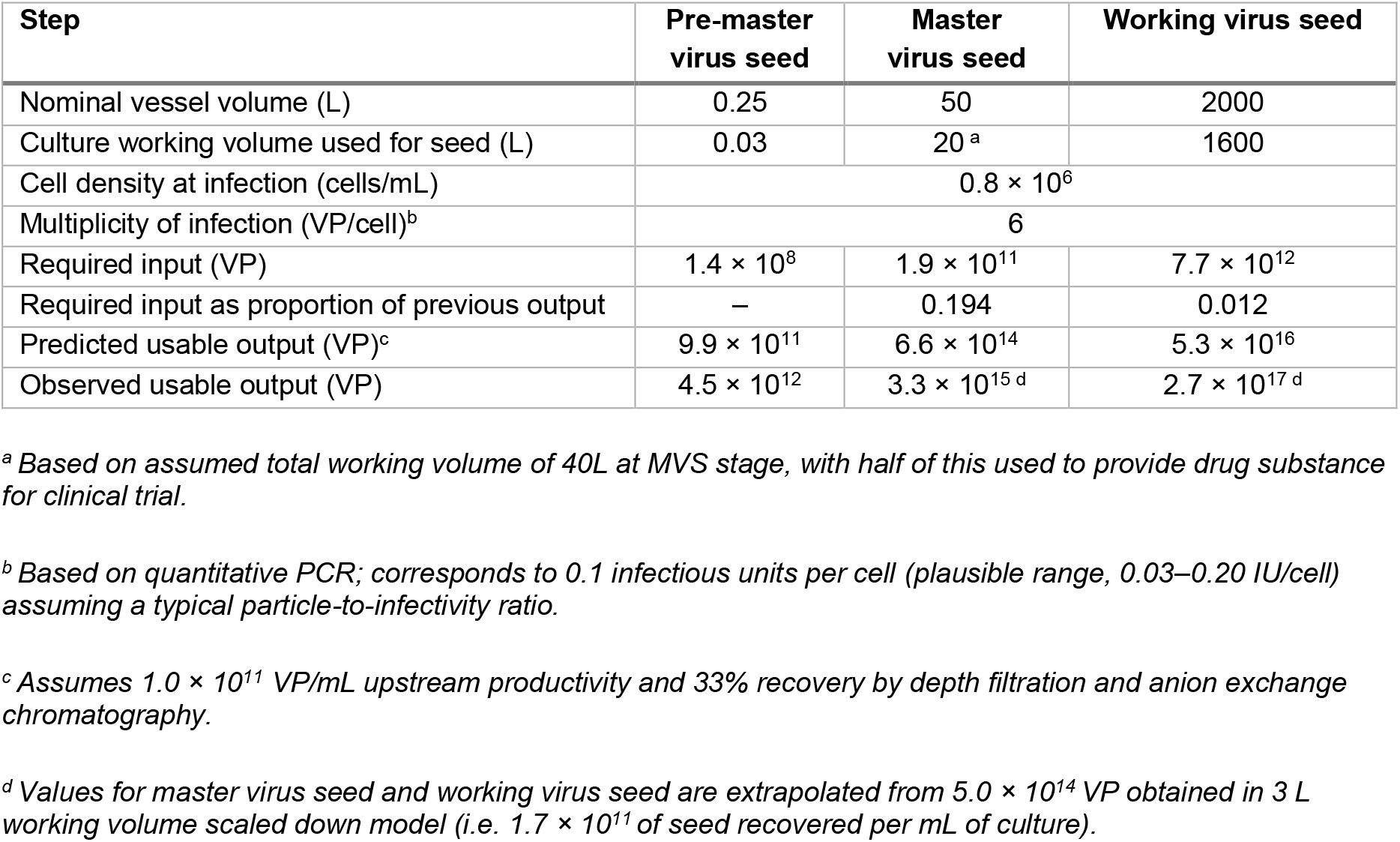
Performance of workflow for working virus seed production.

To test this workflow we performed a simulated low-MOI pre-master working virus seed step at 30 mL scale. After infection at a qPCR-determined MOI of 6 VP/cell, we obtained an output of 4.5 × 10^12^ VP, or greater than 20 times more than required in our workflow to infect a 50 L bioreactor for production of working virus seed. We used a 3 L culture volume in a stirred-tank bioreactor as a model of both 50 L scale master virus seed production and 2000 L working virus seed production. This provided 1.7 × 10^11^ VP/mL of culture, again substantially exceeding that required by the workflow (Table 1).

Using test turnaround times based upon our previous experience, we estimate that this workflow could allow 10 000 doses of vaccine to be released to clinical trial within 60 days of antigen sequencing. Working viral seed could be provided to globally distributed large-scale manufacturing sites at the same time. This in turn could permit release of fully tested vaccine at large scale after a further 35 days, i.e. in a total of less than 100 days from pathogen identification (Figure 1).

### CELL-DENSITY EFFECT IN A MODEL FED-BATCH PROCESS

The above work suggested that relatively simple modifications to our viral seed production process could achieve speed compatible with the CEPI ‘100 day’ target.(1). We next wished to improve volumetric productivity, to increase the rate of vaccine output beyond the first batch.

We initially sought to determine the effect of cell density in limiting volumetric productivity of ChAdOx1 nCoV-19 in T-REx-293 cells, using a scaled-down fed-batch process in shake flasks. Volumetric productivity was not improved by increasing cell density above 4 × 10^6^ cells/mL at infection, due to declining cell-specific productivity (Figure 2). Maximal volumetric productivity before downstream processing was approximately 4 × 10^11^ VP/mL.

**Figure 2.**
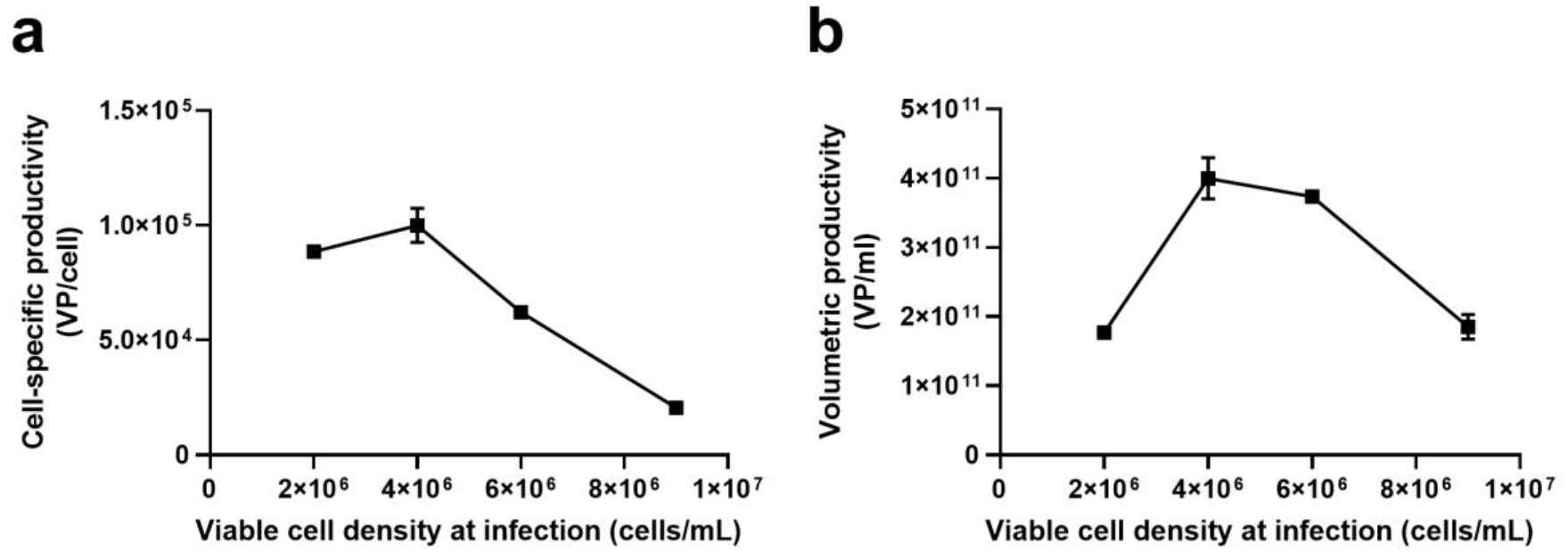
Limitation of productivity by the ‘cell density effect’. (**a**) Cell-specific and (**b**) volumetric productivity of ChAdOx1 nCov-19 over a range of viable cell density at infection. Symbols show median values and error bars show range (if large enough to display) from duplicate shake flasks. Productivity values were based on viral particles measured by qPCR. Viable cell density was measured at infection. Abbreviations: VP, viral particles; qPCR, quantitative PCR.

### HIGH-THROUGHPUT ASSESSMENT OF MEDIUM EXCHANGE

We next sought to explore conditions under which a perfusion-based process might enhance productivity while adding as little complexity as possible to the process. We used the Ambr250 HT perfusion multi-parallel bioreactor system with ATF-mode medium-exchange units (Figure 3a) to assess the effect of three controlled factors on volumetric and cell-specific productivity: viable cell density at infection, perfusion start time, and duration of intensified perfusion after infection (Figure 3b). High and low levels for each factor were assessed in a 2 × 2 × 2 factorial design, with additional center points, in two independent experiments (Figure 3c; Supplementary Table 1). Across both experiments, this approach provided a parameter space encompassing: perfusion starting at a viable cell density (PSVCD) in the range of approximately 1–7 × 10^6^ cells/mL, a viable cell density at infection in the range of 4–18 × 10^6^ cells/mL, and a duration of intensified perfusion after infection of 0, 24 or 48 h (Figure 3 d–e).

**Figure 3.**
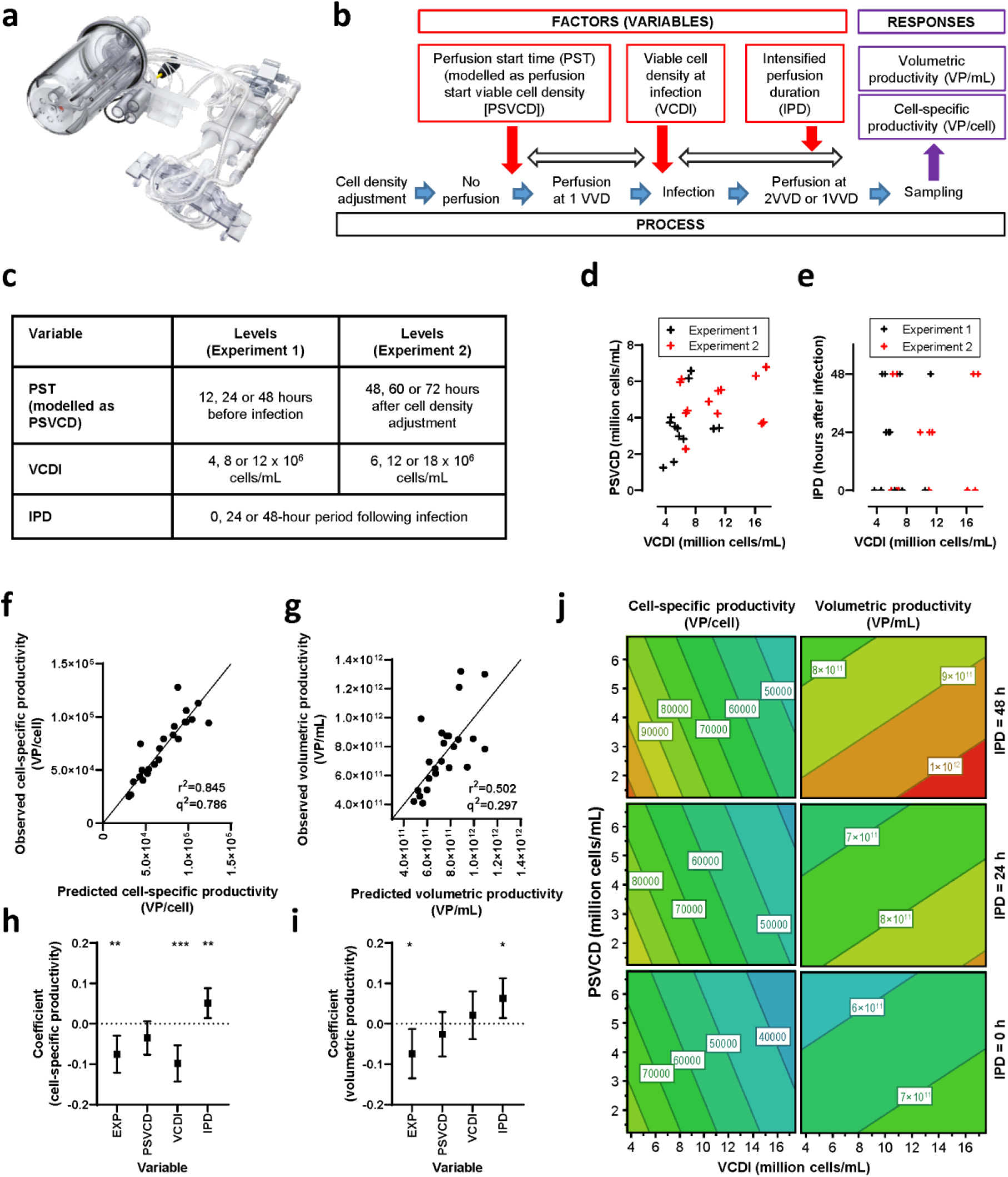
High-throughput assessment of perfusion parameters on productivity. (**a**) Ambr250 perfusion bioreactor system, with perfusion unit for medium exchange via alternating tangential flow filtration. (**b**) Experimental design showing the three factors tested as independent variables and the two responses tested as dependent variables. (**c**) High, low and centre-point levels for each variable in experiment 1 and experiment 2. (**d–e**) Multivariate regression model parameter space for the three variables across each bioreactor in experiment 1 (black) and experiment 2 (red); note that PST is converted to PSVCD for modelling because of the different way in which PST was triggered in the two experiments. (**f–g**) Modelled coefficients for the four factors for each of the two productivity response variables; error bars show 95% confidence intervals. (**h–i**) Scatter plots of observed versus predicted values assessing model goodness of fit for the two productivity response variables; diagonals show lines of identity. (**j**) Contour plots of modelled productivity, with the ‘experiment’ factor held constant at the mean, illustrating relative effects on the responses across the ranges of the variables but not representing predictions; modelled values are likely to be particularly unreliable beyond the experimental design space. *p < 0.05, ** p < 0.01, *** p < 0.001 (t test). Abbreviations: IPD, intensified perfusion duration; PST, perfusion start time; PSVCD, perfusion start viable cell density; VCDI, viable cell density at infection; VP, viral particles; VVD, vessel volume(s) per day.

Following regression modelling, scatter plots of actual versus predicted values showed acceptable model fits, with *r^2^* values of 0.845 for cell-specific productivity and 0.502 for volumetric productivity (Figure 3 f–g). Observed data for each bioreactor are shown in Supplementary Figure 1. Sensitivity analyses indicated that the estimated factor coefficients were robust to a variety of changes in the approach to analysis, including analysis of the first and second experiments in isolation (Supplementary Figure 2).

The regression model indicated that longer intensified perfusion duration significantly improved both cell-specific productivity (*p* < 0.01) and volumetric productivity (*p* < 0.05) (Figure 3 h–i). Lower viable cell density at infection significantly improved cell-specific productivity (*p* < 0.001) but not volumetric productivity. The effect of perfusion start viable cell density was not statistically significant, but trended towards increased productivity with earlier perfusion start. Productivity metrics were also statistically significantly influenced by whether the bioreactor was part of the first or second experiment (which used slightly different cell culture conditions). We did not consider this to be of concern, as our aim was to gather information on the influence of each factor (which was consistent between experiments, as per the sensitivity analyses), rather than to predict absolute values of the responses in a hypothetical future experiment.

Under the optimal modelled conditions, cell-specific productivity before downstream processing was predicted to exceed 10^5^ VP/cell and volumetric productivity to exceed 10^12^ VP/mL. (Figure 3j). The highest observed volumetric productivity was 1.3 × 10^12^ VP/mL, in two bioreactors in the first experiment which had viable cell density at infection of 6 and 11 × 10^6^ cells/mL (Supplementary Table 1). This was not associated with impaired cell-specific productivity, which remained at approximately 1 × 10^5^ VP/cell in both these bioreactors. Volumetric and cell-specific productivity were close to these maximum values in simplified center-point-like bioreactors in both experiments, where perfusion was started early but was not intensified after infection (Supplementary Table 1).

### IMPROVED PERFUSION-BASED UPSTREAM PROCESS

We next tested a new upstream process using ATF medium exchange in three independent accurately simulated production runs in 3L single-use bioreactors. Conditions were similar to those in the simplified center-point-like bioreactors in the multi-parallel experiments, with early perfusion start but no intensification. Perfusion with 1 vessel volume per day starting 48 hours before infection, at a viable cell density of approximately 3 × 10^6^ cells/mL, resulted in a viable cell density at infection of 6.5–7.0 × 10^6^ cells/mL across the three runs (Figure 4a; Table 2). Viable cell density peaked at around 1 × 10^7^ cells/mL and viability then declined as expected during viral replication.

**Figure 4.**
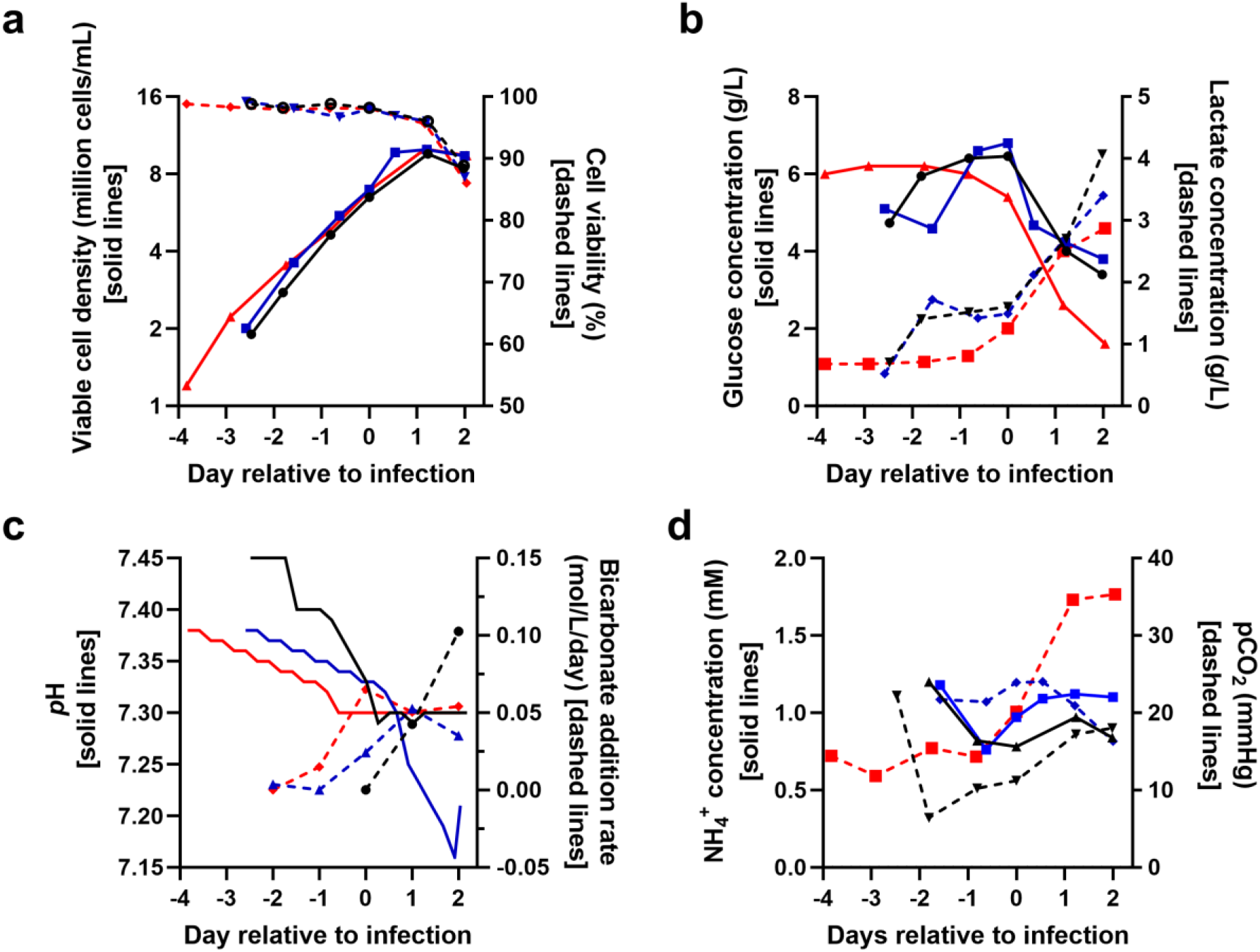
Perfusion-based upstream process performance in three independent scaled-down production runs. Data from three independent production runs conducted on separate occasions; experiment referred to as CJ74 in black, CJ78 in red and CJ87 in blue. Medium exchange via ATF perfusion started at one vessel volume per day at day −2 relative to infection and continued to day +2 relative to infection, when the bioreactors were harvested. (**a**) Viable cell density and cell viability. (**b**) Glucose and lactate concentration. (**c**) pH and rate of bicarbonate addition. (**d**) ammonia concentration and partial pressure of carbon dioxide. See Table 2 for productivity data.

**Table 2.** Productivity and performance of the improved perfusion-based upstream process. Table 2 shows data from the three experiments reported in Figures 4 and 5. Further information about downstream process performance is provided in Supplementary Table 2.

| Parameter | CJ74 | CJ79 | CJ87 |
| --- | --- | --- | --- |
| **Upstream process** |  |  |  |
| Volumetric productivity (VP/mL of culture) | $1.4 \times 10^{12}$ | $1.6 \times 10^{12}$ | $1.6 \times 10^{12}$ |
| Viable cell density at start of perfusion (cells/mL) | $2.8 \times 10^6$ | $3.6 \times 10^6$ | $3.5 \times 10^6$ |
| Viable cell density at infection (cells/mL) | $6.5 \times 10^6$ | $7.0 \times 10^6$ | $6.9 \times 10^6$ |
| Viable cell density at peak (cells/mL) | $9.6 \times 10^6$ | $1.0 \times 10^7$ | $1.0 \times 10^7$ |
| Cell-specific productivity (VP/cell) based on peak viable cell density | $1.5 \times 10^5$ | $1.6 \times 10^5$ | $1.6 \times 10^5$ |
| **Complete process** |  |  |  |
| Volumetric productivity (VP of DS / mL of upstream culture) | $8.1 \times 10^{11}$ | $8.3 \times 10^{11}$ | $7.9 \times 10^{11}$ |
| Particle:infectivity ratio | 35 | 42 | 77 |
| Host cell protein (ng/mL) | 33 | 61 | 79 |
| A260:A280 ratio | 1.17 | 1.18 | 1.15 |
| Host cell DNA (ng/dose) | <0.1 | <0.1 | <0.1 |

Glucose concentrations decreased and lactate concentrations increased from the day of infection onwards (Figure 4b). Small ongoing reductions in *p*H were associated with increasing rates of bicarbonate addition from the day of infection onwards (Figure 4c), with little change in ammonia and carbon dioxide levels (Figure 4c). ATF filter transmembrane pressure remained stable between −0.8 and −0.9 bar throughout perfusion, indicating no significant fouling of the membrane.

Volumetric productivity before downstream processing was 1.4–1.6 × 10^12^ VP/mL across the three simulated production runs and cell-specific productivity was 1.5–1.6 × 10^5^ VP/cell (Table 2).

### COMPATIBILITY WITH EXISTING DOWNSTREAM PROCESS

To assess compatibility of our new upstream perfusion-based process with the existing downstream process, we purified product from the three independent upstream runs using two versions of our previously published method (5). These comprised clarification by combined depth filtration and 0.2 μm filtration (Figure 5a), purification by anion-exchange chromatography (Figure 5b), and formulation by tangential-flow filtration and 0.2 μm filtration. Product recovery after each step and filter loadings were within the expected ranges (Supplementary Table 2). Addition of an extra tangential-flow filtration step after clarification in one run did not alter product recovery (Supplementary Table 2).

**Figure 5.**
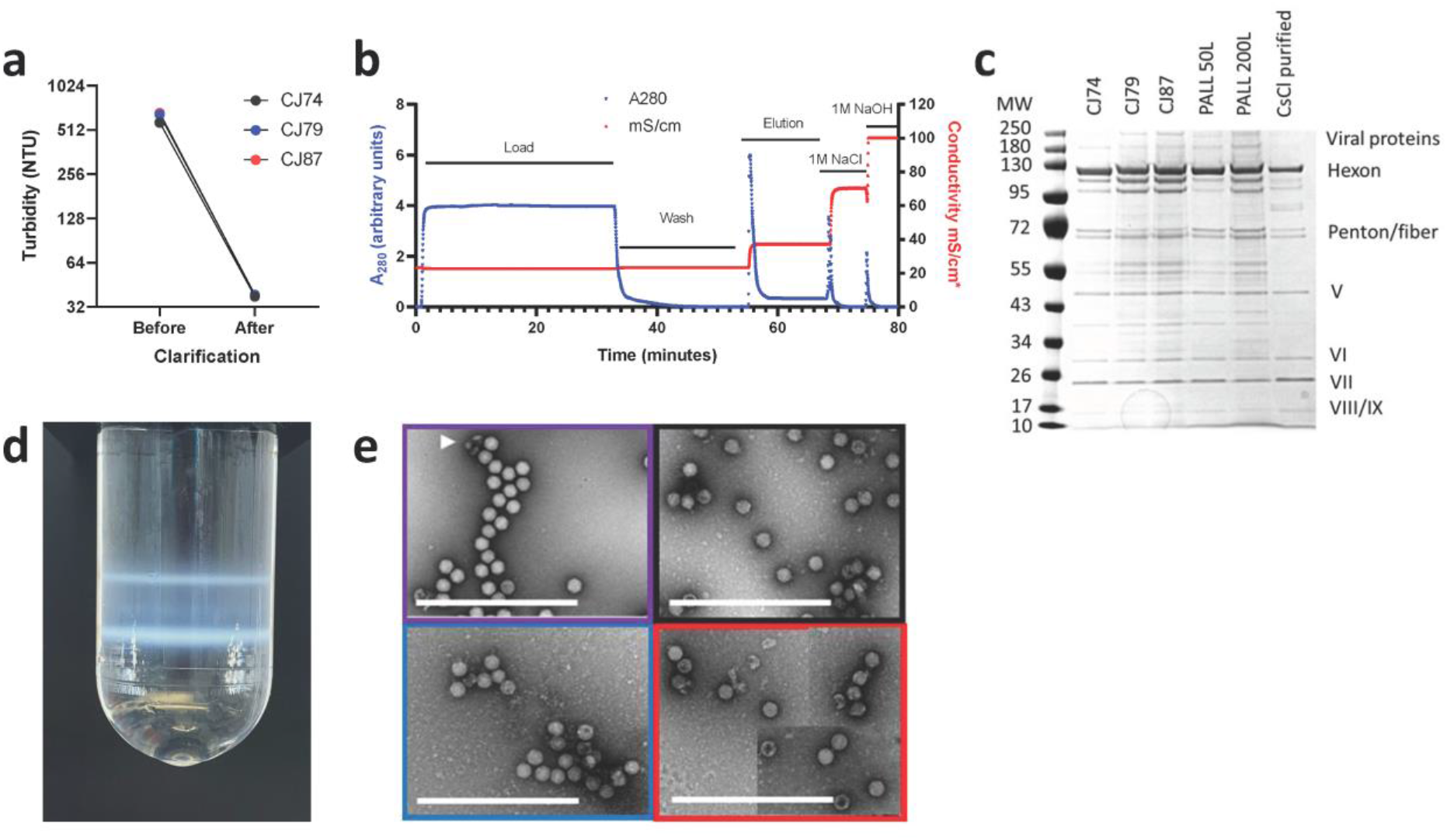
Downstream process performance and product characterization. **(a)** Turbidity before and after clarification of crude cell lysate from the three independent model production runs conducted on separate occasions: CJ74 in black, CJ78 in red and CJ87 in blue. **(b)** Representative anion exchange chromatogram for purification of product from these three runs. Note that maximal measurable conductivity was 100 mS/cm. **(c)** Silver-stained denaturing and reducing sodium dodecyl sulphate polyacrylamide electrophoresis gel loaded with downstream processed material from the three model production plus two controls from actual GMP manufacturing runs (PALL) and CsCl density gradient ultracentrifugation-purified product. Note that all production samples contain additional bands, some of which will correspond to empty viral capsids compared with CsCl-purified control, as expected for the chromatographic downstream process. **(d)** Representative analytical CsCl density gradient ultracentrifugation from one of the three model production runs showing upper band corresponding to empty viral capsid. **(e)** Representative negative-stain transmission electron micrographs of product from the three model production runs, colour coded as in panel a, together with CsCl ultracentrifugation-purified comparator (purple). Note that empty capsids admit and complete capsids exclude the contrast agent. Arrowhead indicates artefactual fragments resulting from sample preparation.Scale bars indicate 1μm. Red-bordered image is a composite of four separate images.

Quality assays indicated that particle-to-infectivity ratio and host cell protein and DNA content were within the acceptable ranges for all three runs (Table 2). The product contained the same viral capsid proteins as product derived from the conventional process (Figure 5c). The presence of an expected minor component of empty viral capsids, consistent with chromatographic purification, was confirmed by density gradient ultracentrifugation (Figure 5d) and electron microscopy (Figure 5e). Volumetric productivity after downstream processing was approximately 8 × 10^11^ VP of drug substance per mL of bioreactor culture, with very similar values across the three runs (Table 2), representing downstream efficiency of approximately 50%. This would be sufficient for >10 000 finished doses per L of bioreactor working volume.

This productivity implies that, despite the increased seed requirements of a high MOI process, WVS from a single 2000L bioreactor run (table 1) would be sufficient to produce >100m finished doses. This calculation is based upon assumptions we believe to be cautious i.e. output of 5.3×10^16^ VP per 2000L WVS reactor, particle: infectious unit ratio of 100, 33% wastage due to QC and aliquoting, and infection at 6 × 10^6^ cells/mL with an MOI of 5 IU/cell.

### TECHNO-ECONOMIC MODELLING

We next constructed a techno-economic model using Biosolve software, informed by our small-scale experimental data and our previous model of a 2000 L fed-batch process (5) to estimate the costs, consumables and equipment requirements for a 2000L perfusion-based process.

We examined two scenarios for use of the seed train, production bioreactor and downstream process train. The first scenario comprised a single seed train, culminating in a 200 L non-perfusion ‘n–1’ reactor, a single 2000 L production reactor, and a single downstream process train. This required slow turnaround of the production vessel (c. 10 day cycle time, comprising 7 days of cell expansion, 2 days of infection, 1 day of cleaning/ preparation) and hence low utilization of the downstream process train. In the second scenario, a 200 L perfusion-enabled ‘n–1’ reactor was added to the seed train (reducing the duration of cell expansion in the production bioreactor) and used to serve two 2000 L production reactors. This approach would provide a batch for downstream processing approximately every 4 days (Figure 6). By more than doubling utilization of the seed train and purification train, this approach could more than double output from a suite with a proportionally small increase in required footprint. Modelling of the downstream process suggested it could be executed with moderate modification to the equipment used for the fed-batch process, and within the capacity of typical off-the-shelf equipment (Figure 6). The large volumes of buffer required for AEX are notable: we have not attempted to reduce buffer requirements at this stage and believe it is likely to be feasible to achieve substantial reduction by shortening equilibration and wash steps.

**Figure 6.**
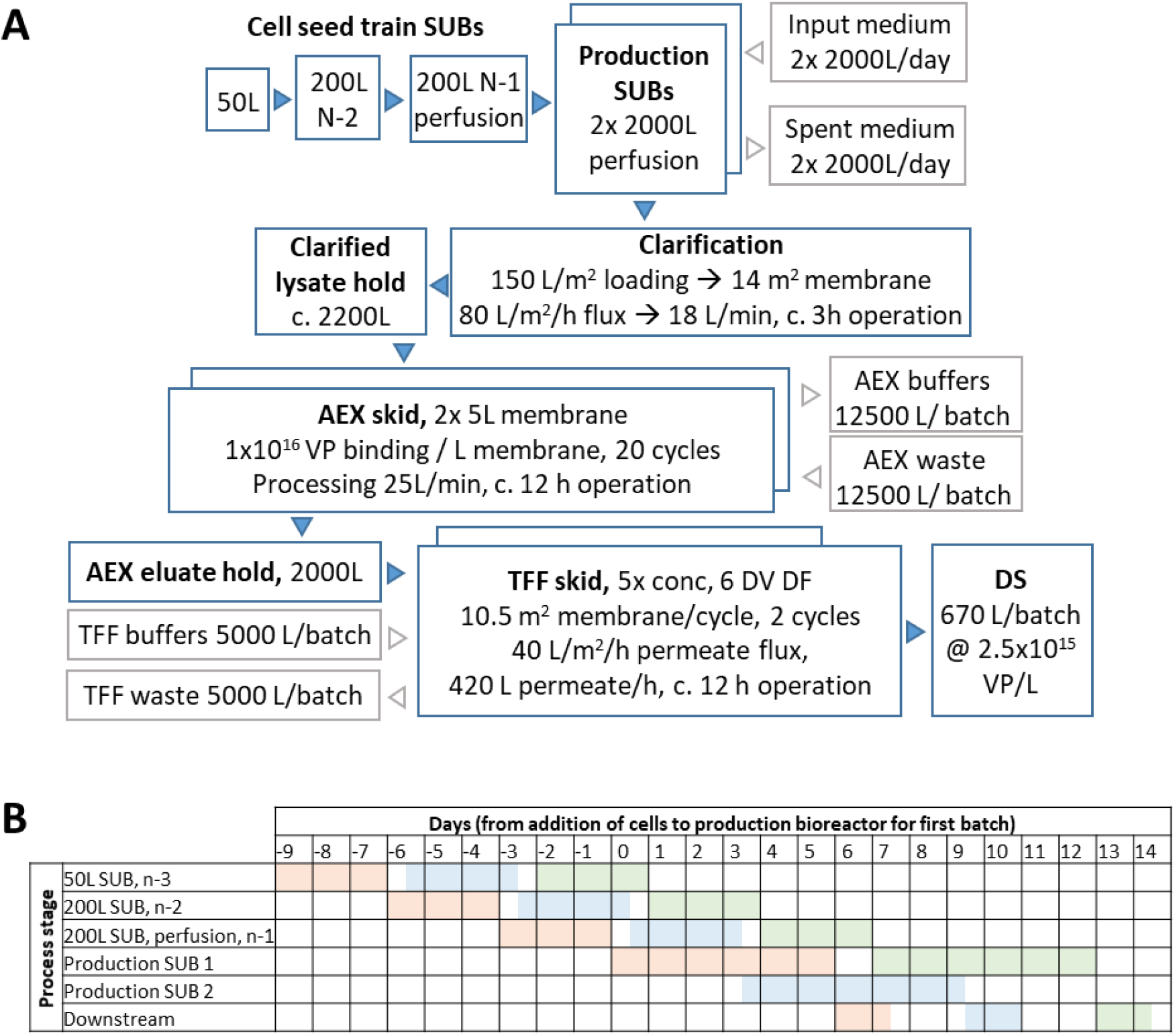
Techno-economic modelling of perfusion process. **(a)** Equipment, product and materials flow in a cleanroom equipped as proposed with 2× 2000L perfusion-capable bioreactors. **(b)** Illustrative production schedule, showing use of single cell seed train, two production bioreactors and single downstream purification train. This schedule provides a batch every 3.5 days; calculations of facility output in text assume a more conservative 4 day cycle time. Abbreviations: SUB – single-use bioreactor; AEX – anion exchange; TFF – tangential flow filtration; DS – drug substance.

Assuming 70% facility utilization, a single such facility is predicted to be capable of production of approximately 1.2 billion doses per year, with a cost of goods below 0.20 EUR/dose. Under an alternative assumption of short-term maximum-capacity operation for emergency response, we estimate that 8 such facilities (i.e. total installed bioreactor capacity of 32 000L) could provide a total of one billion doses per month.

## Discussion

The SARS-CoV-2 genome sequence was published on 10 Jan 2020.(19) The adenovirus-vectored vaccine ChAdOx1 nCoV-19 was administered to the first volunteer in a clinical trial after 104 days (23 April 2020) and administered for the first time outside a clinical trial after 360 days (4 January 2021).(20) Release of the billionth dose was announced after 566 days (29 July 2021).(21) The first mRNA vaccine (from Pfizer) and inactivated SARS-CoV-2 vaccine (from Sinovac) to release a billion doses did so around the same time. (22) The rates of progress of leading COVID-19 vaccines to each of these milestones have been substantially faster than had been achieved for any previous novel vaccine.

We now estimate that, if our current understanding had been available immediately at the time of SARS-CoV-2 sequence publication, and production had proceeded as described here, a billion finished doses could have been available for release more than a year earlier, before the end of May 2020. This would have required significant investment ‘at financial risk’, and the availability of both drug substance and drug product production facilities, staff, equipment and materials ‘on standby’. Translation of this manufacturing speed into rapid public health impact would require equally fast pre-clinical and clinical development (the same applies to other vaccine platforms). This may or may not be achievable for a completely new pathogen, but almost certainly is achievable for a new version of an existing vaccine in response to a new pathogen variant (for example, SARS-CoV-2, or potentially influenza).

We expect similar acceleration of manufacturing will also be possible for other vaccine platforms. The factor limiting speed of vaccine availability may be speed of early decision-making. Willingness to invest ‘at financial risk’ in the early stages of seed generation, facility preparation and clinical trial preparation, long before it is even clear there is a public health need for any vaccine in response to a new and incompletely characterized outbreak provides a global public good.

For an adenovirus-vectored vaccine, rapid seed generation remains a key determinant of time to trial-scale and large-scale vaccine supply. Because the seed generation and perfusion processes we describe here are independent of each other, and the seed generation method involves only modest change to existing processes, we believe that the timelines we outline for trial-scale and large-scale supply could be achieved almost immediately. The main changes required are adoption of low-MOI two-cycle processes, which are already well-established for drug substance production, at pre-master and master and working virus seed stages. We note also the need for analytical demonstration of comparability of vaccine produced using different upstream processes for the initial trial batch and large-scale production. Speed of the process depends critically on willingness of the GMP seed production facilities to accept incoming starting material before microbiological testing is completed. We believe that each step of the seed production process is robust and that risks of assay failures are low but it would nonetheless seem prudent, on both public health and financial grounds, to invest additional effort in mitigating risks of failure. A fully standard-operating-procedure-defined and validated version of the process described here, preferably including an independent parallel ‘back up campaign’ and supported by rapid-turnaround platform analytics (notably easily-accessible nucleic-acid based adventitious agent assays), would in our view provide high confidence of success. Beyond first large-scale supply, the time to reach output milestones (such as the billionth dose) is a function of the resources committed (e.g. m^2^ of production facility) and the output in doses per m^2^ per day. The latter is a function of volumetric productivity and cycle time. The novel perfusion-based upstream process for ChAdOx1 nCoV-19 production which we have described here boosts volumetric productivity by around four fold compared with the current process (reaching approximately 8 × 10^14^ VP of purified drug substance per liter of bioreactor culture, sufficient for >10,000 doses of DP). To our knowledge this is the first demonstration that perfusion can improve productivity of a species E adenoviral vector or a simian adenoviral vector. We are not aware of any other reported process achieving greater productivity with such vectors.

Our ‘design of experiments’ approach on the Sartorius Ambr 250 HT perfusion system identified early perfusion start and intensified perfusion after infection as factors that improved volumetric productivity. We included early perfusion start but not intensified perfusion after infection in the design of a new perfusion-based upstream process for testing in a 3 L system. In this work we demonstrated reproducible performance of the process, using a constant rate of perfusion (1 vessel volume per day) from 48 hours before infection to harvest at 48 hours after infection. Our rationale for not intensifying perfusion was that facilities manufacturing ChAdOx1 nCoV-19 are already capable of handling 1 vessel volume per day of medium (as a requirement of the current process), but may not be capable of handling intensification to 2 vessel volumes per day. The large improvement in upstream productivity is compatible with the existing downstream process. We demonstrated product recovery (approximately 50%) and product quality in line with expectations.

The work described here was facilitated by the fact that adenovirus production remains a brief batch process, rather than a long continuous process (in which issues such as filter fouling may be more problematic). The work was also facilitated by use of novel scaled down model systems. We believe this may be the first published report of such multi-parallel optimization of any perfusion process (‘design of experiments’ in a total of 25 reactors, enabled by the first published use of the Ambr250 HT perfusion system). It may also be the first published report of use of the XCell Lab ATF controller and ATF1 filter system (enabling accurate modelling of ATF filter dynamics in a 2000 L process at 3 L laboratory scale).

We used a techno-economic model to consider equipment/facility requirements and campaign design to enable high output (doses per m^2^ per day) from this process. We estimate that a single facility housing two 2000L production bioreactors might achieve short-term output of around 120m doses/month. We believe numerous existing facilities (including several in low-and middle-income countries) would be suitable for this. Cost per dose of drug substance would be expected to be around half that of that achieved with the current fed-batch process. There is clearly uncertainty in making projections based upon linear scale-up of a small-scale process, but our 3L process was designed carefully as an accurate scale-down model of 2000L-scale production, we and others have previously had reasonable success in scale-up of similar processes, and the downstream process here is essentially unchanged from that already in use.

Overall, this work suggests suitability of the adenovirus-vector vaccine platform to hit the CEPI ‘100-day’ moonshot goal, and feasibility of considerable improvement upon the productivity of a manufacturing process which already, at the time of writing, is estimated to have delivered more COVID-19 vaccine doses than any other.(23) We believe there remains scope for further improvement in the time to both trial-scale and large-scale supply, and the facility footprint required to achieve a given level of output. The methods described here, however, are likely to be sufficient to reduce differences in manufacturing speed and facility output versus other platforms, including mRNA, to a level at which other factors become more important in considering the ongoing roles of different platforms in emergency response. Such considerations include safety, tolerability, efficacy, stability, cost, programmatic suitability, individual recipient preference, and manufacturer willingness to support technology transfer beyond their own facilities. With respect to these other factors, the adenovirus platform has strengths which are currently unique, complementary to alternatives, and in some cases are context-specific.

Public health benefit is likely to be maximized by availability of diverse vaccines, including adenoviruses, and there may be valid scientific reasons for selection of different vaccines in different contexts. Emerging evidence suggests durable protection with slower decay of antibody responses and superior CD8^+^ T cell immunogenicity after single-dose adenovirus-vectored vaccination, as compared to two doses of mRNA.(24–26) Thrombosis with thrombocytopenia syndrome, which has occurred very rarely in certain populations given adenovirus-vectored vaccines, appears even rarer in studied data sets from outside Europe and North America.(27–29) Accumulation of safety experience comparable to that of the most widely used COVID-19 vaccines – including adenovirus, mRNA and inactivated virus platforms – will take significant time for new vaccines based on alternative platforms. Important and continuing advantages of adenovirus-vectored vaccines are the existence of a global network of manufacturing facilities, well-characterised safety, ability to achieve single-dose protection lasting months, and the ease of distribution enabled by ≥6 month stability at 2-8 °C in liquid formulations.(30, 31)

## Acknowledgements

The authors are grateful for the assistance of Adam Ritchie, the Viral Vector Core Facility at the Jenner Institute, University of Oxford, and Errin Johnson from the electron microscopy core facility at the Sir William Dunn School of Pathology, University of Oxford. Sarah Gilbert and Teresa Lambe designed the ChAdOx1 nCoV-19 vaccine. We also wish to thank Richard Turner, Raghavan Venkat, Jinlin Jiang, Nicole Bleckwenn and Thomas Linke (all at AstraZeneca), and Shelly Parra and Earl Pineda (at Repligen) for helpful discussions.

Under the direction of the authors, Dr M Cottingham from Oxford PharmaGenesis provided medical writing assistance with funding from the University of Oxford.

## Supplement

### SUPPLEMENTARY TABLES

**Supplementary Table 1.** Variable and response levels in each bioreactor.

| Bioreactor | Variables |  |  |  | Responses |  |
| --- | --- | --- | --- | --- | --- | --- |
|  | PDBI<br>(hours) | PSVCD<br>(10<sup>6</sup> cells/mL) | VCDI<br>(10<sup>6</sup> cells/mL) | IPD<br>(hours) | Volumetric<br>productivity<br>(VP/mL) | Cell-specific<br>productivity<br>(VP/cell)<sup>a</sup> |
| **Experiment 1** |  |  |  |  |  |  |
| 1 | 12 | 3.74 | 4.57 | 0 | $8.9 \times 10^{11}$ | $1.3 \times 10^5$ |
| 2 | 48 | 1.25 | 3.67 | 0 | $6.5 \times 10^{11}$ | $9.8 \times 10^4$ |
| 3 | 12 | 6.58 | 7.39 | 0 | $6.2 \times 10^{11}$ | $6.0 \times 10^4$ |
| 4 | 48 | 3.40 | 10.44 | 0 | $8.7 \times 10^{11}$ | $6.0 \times 10^4$ |
| 5 | 12 | 4.02 | 4.71 | 48 | $8.6 \times 10^{11}$ | $1.1 \times 10^5$ |
| 6 | 48 | 1.57 | 5.09 | 48 | $7.8 \times 10^{11}$ | $9.4 \times 10^4$ |
| 7 | 12 | 6.16 | 7.10 | 48 | $6.6 \times 10^{11}$ | $7.9 \times 10^4$ |
| 8 | 48 | 3.43 | 11.19 | 48 | $1.3 \times 10^{12}$ | $8.3 \times 10^4$ |
| 9<sup>b</sup> | 30 | 2.99 | 5.78 | 24 | $1.3 \times 10^{12}$ | $1.1 \times 10^5$ |
| 10<sup>b,d</sup> | 30 | 3.52 | 5.24 | 24 | $8.5 \times 10^{11}$ | $9.5 \times 10^4$ |
| 11<sup>b</sup> | 30 | 3.42 | 5.59 | 24 | $1.2 \times 10^{12}$ | $9.5 \times 10^4$ |
| 12<sup>c</sup> | 48 | 2.83 | 6.40 | 0 | $8.7 \times 10^{11}$ | $9.1 \times 10^4$ |
| **Experiment 2** |  |  |  |  |  |  |
| 1 | 24 | 4.39 | 6.93 | 0 | $5.0 \times 10^{11}$ | $5.1 \times 10^4$ |
| 2 | 0 | 5.95 | 5.95 | 0 | $4.2 \times 10^{11}$ | $4.7 \times 10^4$ |
| 3 | 72 | 3.76 | 17.15 | 0 | $5.0 \times 10^{11}$ | $2.7 \times 10^4$ |
| 4 | 48 | 6.30 | 16.09 | 0 | $4.6 \times 10^{11}$ | $2.5 \times 10^4$ |
| 5 | 24 | 4.25 | 6.71 | 48 | $7.0 \times 10^{11}$ | $7.9 \times 10^4$ |
| 6 | 0 | 6.13 | 6.13 | 48 | $6.5 \times 10^{11}$ | $7.0 \times 10^4$ |
| 7 | 72 | 3.68 | 16.90 | 48 | $8.0 \times 10^{11}$ | $4.4 \times 10^4$ |
| 8 | 48 | 6.79 | 17.50 | 48 | $8.2 \times 10^{11}$ | $3.9 \times 10^4$ |
| 9<sup>b,e</sup> | 36 | 4.88 | 9.79 | 24 | $6.2 \times 10^{10}$ | $5.8 \times 10^3$ |
| 10<sup>b</sup> | 36 | 5.47 | 11.03 | 24 | $5.8 \times 10^{11}$ | $4.0 \times 10^4$ |
| 11<sup>b</sup> | 36 | 5.53 | 11.43 | 24 | $6.9 \times 10^{11}$ | $5.0 \times 10^4$ |
| 12<sup>c</sup> | 48 | 4.23 | 10.96 | 0 | $9.9 \times 10^{11}$ | $7.5 \times 10^4$ |
| 13<sup>c,d</sup> | 48 | 2.28 | 6.68 | 0 | $4.1 \times 10^{11}$ | $5.5 \times 10^4$ |

<sup>a</sup> Based on peak viable cell density (not shown)
<sup>b</sup> Center-point condition
<sup>c</sup> Simplified center-point-like condition
<sup>d</sup> Outlier with low cell density
<sup>e</sup> Contaminated and excluded from analyses
PDBI, perfusion duration before infection; PSVCD, perfusion start viable cell density; VCDI, viable cell density at infection; IPD, intensified perfusion duration; VP, viral particles.

**Supplementary Table 2.** Step recovery and filter loading in three independent downstream process runs

| PARAMETER | EXPERIMENT |  |  |  |
| --- | --- | --- | --- | --- |
|  | CJ74 | CJ79 | CJ87 | Mean |
| **Step recovery by qPCR** |  |  |  |  |
| Clarification | 73% | 63% | 63% | 66% |
| Pre-AEX TFF* | 95% | N/A | N/A | N/A |
| AEX | 89% | 100% | 83% | 91% |
| Formulation TFF | 98% | 82% | 88% | 89% |
| 0.2 µm filtration | 92% | 92% | 100% | 95% |
| Cumulative recovery (DS as % of upstream) | 56% | 48% | 46% | 50% |
| **Filter loadings** |  |  |  |  |
| Clarification (cells/m<sup>2</sup>) | $1.1 \times 10^{12}$ | $1.1 \times 10^{12}$ | $1.1 \times 10^{12}$ | |
| AEX (VP per mL of membrane) | $6.3 \times 10^{12}$ | $8.2 \times 10^{12}$ | $1.2 \times 10^{13}$ | |

\*Additional step used in CJ74 only.
Abbreviations: AEX, anion-exchange chromatography; TFF, tangential-flow filtration; VP, viral particles.

### SUPPLEMENTARY FIGURES

**Supplementary Figure 1.**
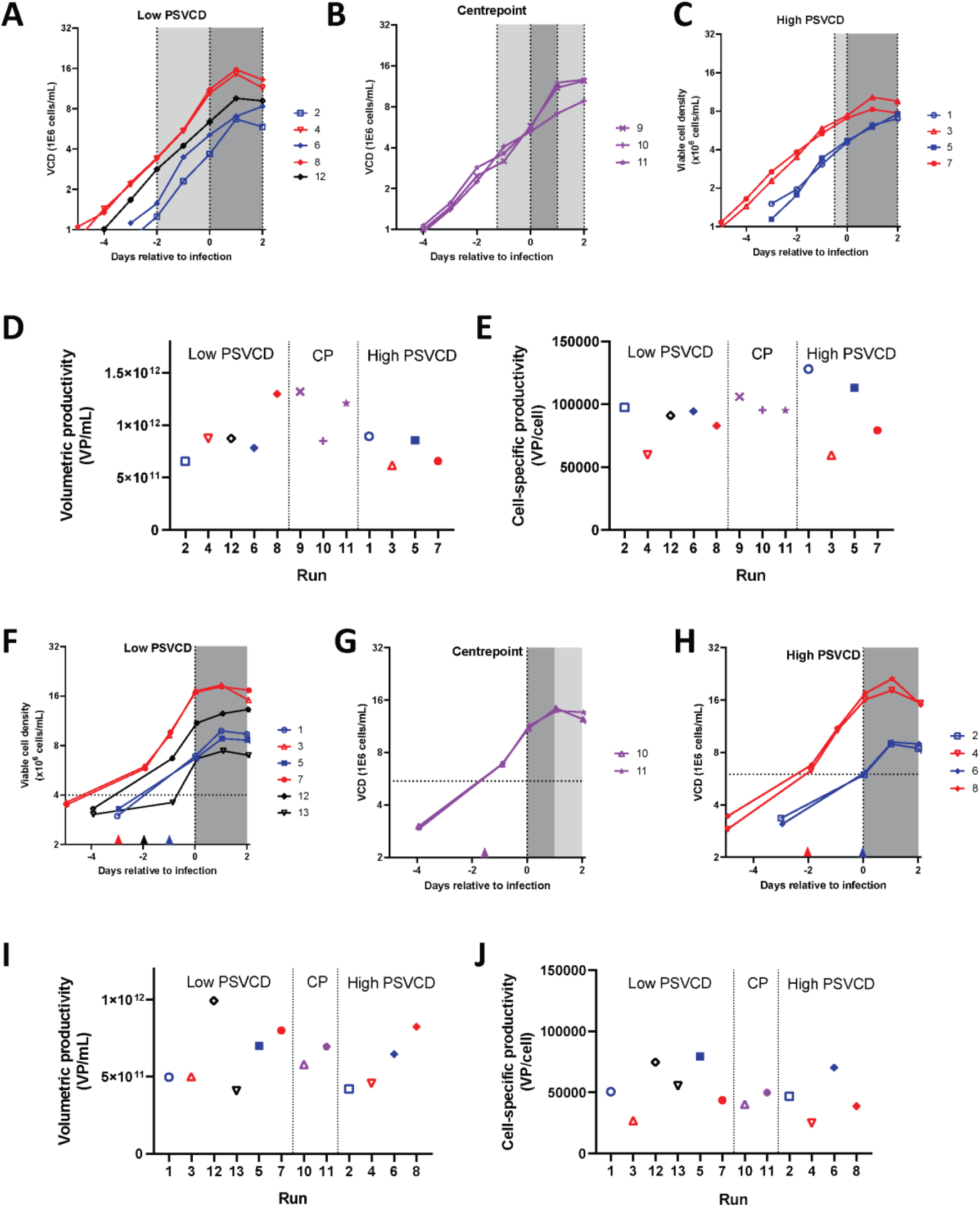
Cell growth and productivity in multi-parallel scaled-down bioreactor experiments. Data in panels A–E is from Experiment 1; data in panels F–J is from experiment 2. Panels A–C and F–H show cell growth in each reactor; D–E and I–J show response variables. Filled symbols indicate intensified perfusion for 48 h after infection; open symbols indicate no intensified perfusion (NB, center point reactors used intensified perfusion for 24 h after infection). Blue indicates low and red indicates high viable cell density at infection; purple indicates center point and black indicates simplified centerpoint like viable cell density at infection. Light grey shading (A–C; G) or arrowheads (F–H) indicate non-intensified perfusion. Dark grey shading indicates intensified perfusion in some reactors. Abbreviations: CP, center point; PSVCD, perfusion start viable cell density.

**Supplementary Figure 2.**
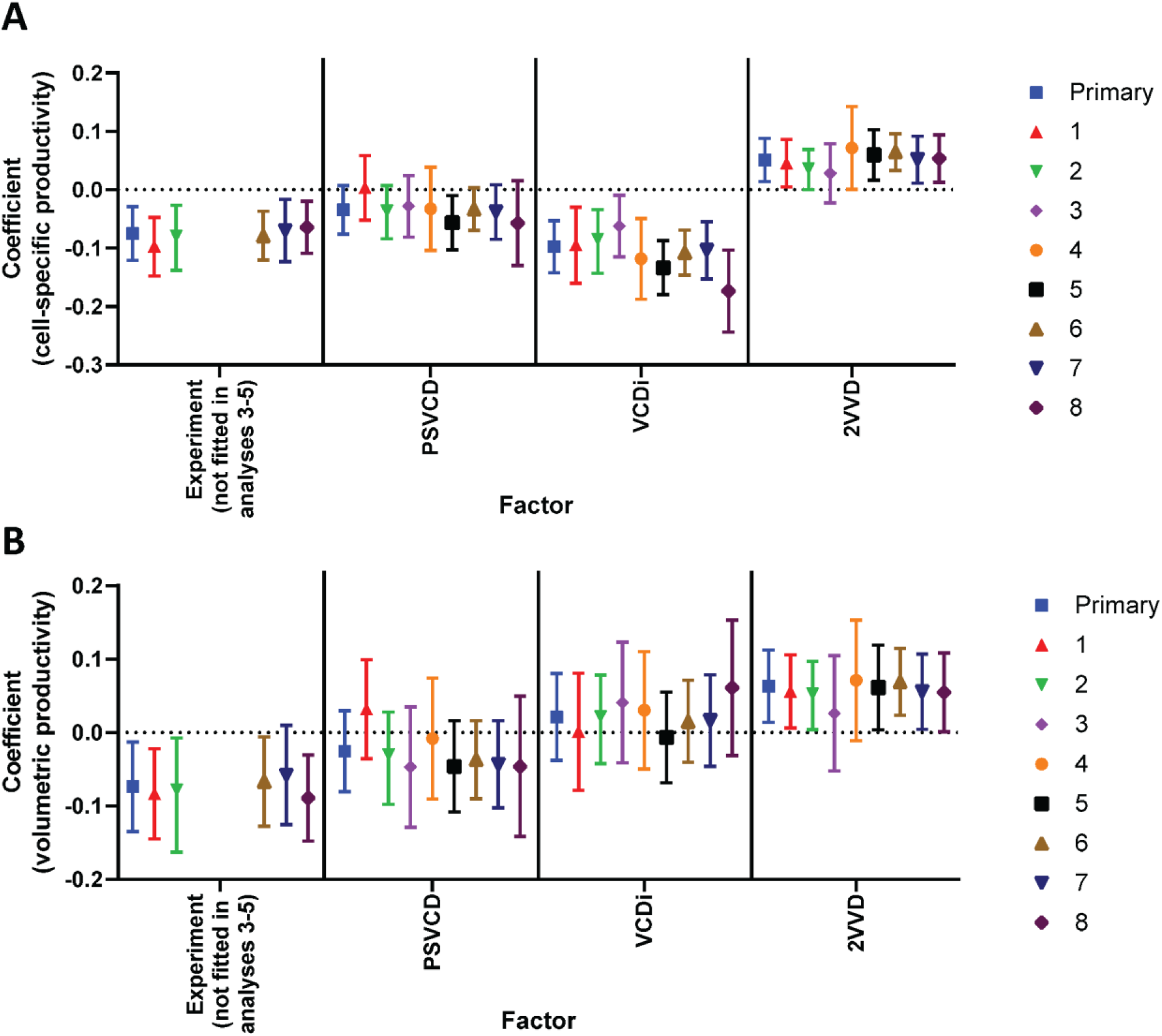
Sensitivity analyses of multi-parallel scaled-down bioreactor experiments for (A) cell-specific productivity and (B) volumetric productivity. Legend: 1, PST factor replaced with perfusion duration before infection (PDBI) instead of PSVCT; 2, raw response data instead of log_10_-transformed data; 3–4, experiments analysed individually instead of in combination; 5, pooling of experiments without the EXP dummy variable; 6, exclusion of three bioreactors with simplified centre-point-like conditions that did not fit the full-factorial design; 7, exclusion of two bioreactors due to initial low cell density likely resulting from error during set-up (one centre-point reactor in the first experiment and one simplified centre-point-like reactor in the second experiment); 8, model fit using multiple linear regression instead of partial least-squares regression. PSVCD, perfusion start viable cell density; VCDi, viable cell density at infection, 2VVD, intensified perfusion with two vessel volumes per day.

